# Invitro-Propagation of Threatened African Sandalwood (Osyris lanceolata Hochst. & Steud.) via Using IBA hormone

**DOI:** 10.1101/2020.01.15.908566

**Authors:** Adugnaw Admas, Semegnwe Melese, Amare Genetu, Berhane Kidane, Zewdu Yilma, Melaku Admasu, Tesaka Misiga

## Abstract

The African Sandalwood plant (Osyris lanceolata) is a threatened shrub or a small hemi-parasitic tree endemic to East Africa and South African regions, which is being severely affected by fungus, uproot-ing for oil extraction, poor natural regeneration, phenological structures (dioecious), medicinal values, lack of sexual recruitment, habitat loss, anthropogenic and climate factors and life span of its seed is short. It has been found a chllange through application of in situ conservation of natural trees like osyris lanceoleta since rapid human population growth and dramatically change the demands of fuel wood and agriculture investement, the available natural strands of valuable plants of African sandalwood have not been able to meet the demands of the people in world specifically developing countries. How ever, this study via using the advances of plant propagation method it provide new options for conserving and multiplication of Osyris lanceolata species using in vitro culture techniques of using IBA hormone by taking plant material of the targated plant.

Propagation of African sandalwood (Osyris lanceolata) by rooting hormone of IBA in non-mist poly-propagator was investigated by taking young stem of osyris lanceoleta from Bazawit Hill and we provide an alternative propagation technique to the use of seeds. This Hill is near to Bahrdar town, Ethiopia.New leaf were initiated on the young stems of the osyris after six weeks of the experment.The influence of IBA as rooting promoter at three concentrations (50, 100 and 150 ppm) were recorded.Un expectedly, from the data collected it was observed that the success 83.3% can be achieved from young stem of un treatead cuting gave a new leaf and 50% were achived in the treatead plant, this propagation technique is a viable alternative to seed. The success may influenced by application orgin of a stem cutting with a type of soil.

## 1. Introduction

Osyris lanceolata is is a shrub or small deciduous tree that grows to 1-7 m in height depending on the soil type, climatic conditions and genetic variation, and has a wide geographic distribution in Africa from Algeria to Ethiopia and south to South Africa, Europe (Iberian peninsula and Balearic Islands), Asia (India to China), and Socotra (Teklehaimanot et al., 2004; Giathi et al., 2011; Kamondo et al., 2012; Gathara et al., 2014). O. lanceolata is distributed in African countries such as Tanzania and Kenya frequently found in arid to semiarid areas, primarily on stony and rocky soils (Kokwaro, 2009) or occasionally in rocky sites and along the margins of dry forests, evergreen bushland, grassland, and thickets at an altitude range of 900-2250 m above sea level (Giathi et al., 2011; Kamondo et al., 2012).

In East African countries, O. lanceolata constituted an important source of medicine (Mwang’ingo et al., 2010). A decoction of the bark and root is considered to be useful for treating diarrhoea, gonorrhea, chronic mucus infections, and urinary diseases (Teklehaimanot et al., 2004; Kokwaro, 2009), a decoction of the bark in boiling water is used to treat candidiasis and related fungal infections (Masevhe et al., 2015) while the essential oil extracted from the bark is used to treat diarrhoea, chest problems, and joint pains. Fibers from the roots are used in basket making while the strong red dye from the bark and roots is used in skin tanning (Mbuya et al., 1994). Since O. lanceolatais an evergreen tree with long flowering periods, it is a good forage plant (Fichtl and Adi, 1994). The utilization of O. lanceolata in the perfumery and fragrance industries in the early 1900s followed a decline in the resource base of Indian sandalwood (Santalum albumL.) (Mbuya et al., 1994). Several communities in Kenya also use O. lanceolata to produce dyes, to treat various ailments, and to brew herbal tea(Kamondo et al., 2012).

How ever, Osyris lanceolata is critically endangred since propagation of by seeds is difficult due to a limited supply and availability of seed at the right time (being a dioecious species, the spatial distribution of trees affects the reproductive outcome; Mwang’ingo et al., 2008), storage difficulties and thus poor germination (Mbuya et al., 1994). Consequently, several interventional measures are required to conserve O. lanceolata. A study by Mwang’ingo et al. (2004) on the storage and pre-sowing treatments on seed germination demonstrated that the testa covering the embryo plays a significant role in limiting germination by restricting gas and water entry and also acts as a mechanical barrier to embryo growth. However, complete removal of the testa and soaking the zygotic embryo in hot water enhanced seed germination by 66.5%, shortened the time to seedling emergence and promoted early seedling growth (Mwang’ingo et al., 2004). Stem cuttings (8-10 cm long with 3-4 leaves from young trees or seedlings) could be induced to root with a maximum of 15% rooting when dipped first in a fungicide (Bavistin) for 5 min, then in 1% indole-3-butyric acid (IBA) for 6 h, but “this concentration could be increased to 32.5% when 75% of the original leaves were left intact” (Giathi et al., 2011). In the same study (Giathi et al., 2011), the choice of substrate was shown to affect the rooting ability of cuttings, with 30% of cuttings rooting in vermiculite, which was superior to sand, vermiculite + sand (1:1), activated coconut peat and peat. Teklehaimanot et al. (2004) used 50-150 mg/L IBA to enhance root production in young stem cuttings collected in early spring. Mwang’ingo et al. (2006) initiated air layers that were left on parent trees for eight weeks and watered every two days to allow root initiation with the help of three concentrations (50, 100 and 150 mg/L) of IBA during February, June, September, and December: 50 mg/L IBA was optimum for root initiation and June to September was best for air layering with about 80% rooting success after potting plants in sand, forest soil and animal manure (2:1:1) and fertilizing with 5 g/container of NPK (nitrogen: phosphorus, potassium) fertilizer. Machua et al. (2008) achieved 60% rooting success through air layering.

In order to restore the previous stocks of san-dalwood species in its natural stands, conventional breeding of sandalwood for introgression of new genetic in-formation can be used. However, it is an expensive and difficult task because of its long generation time, sexual incompatibility and heterozygous nature (Rugkhla and Jones, 1998). Therefore, this research was attempted to addreses the endangred sandlwood via the approaches of using rooting hormone propagation techiniqes in non-mist poly-propagator.

## 2. Material and Method

### 2.1. Study site description

Stem cuttings were collected in Novmbere, 2019 at Bezawit Hill, Bahrdar town. It is far two-and-a-half kilometres south of from the Martyrs Memorial, Bahrdar town, Northen Ethiopia.It was palace of Haile Selassie.The experment were conducted at Bahrdaer plant tissue culture laboratoery.It is near to Bazawit hill. The Bazawit Hill vegetation cover is predominantly grass.

**Figure 1.**
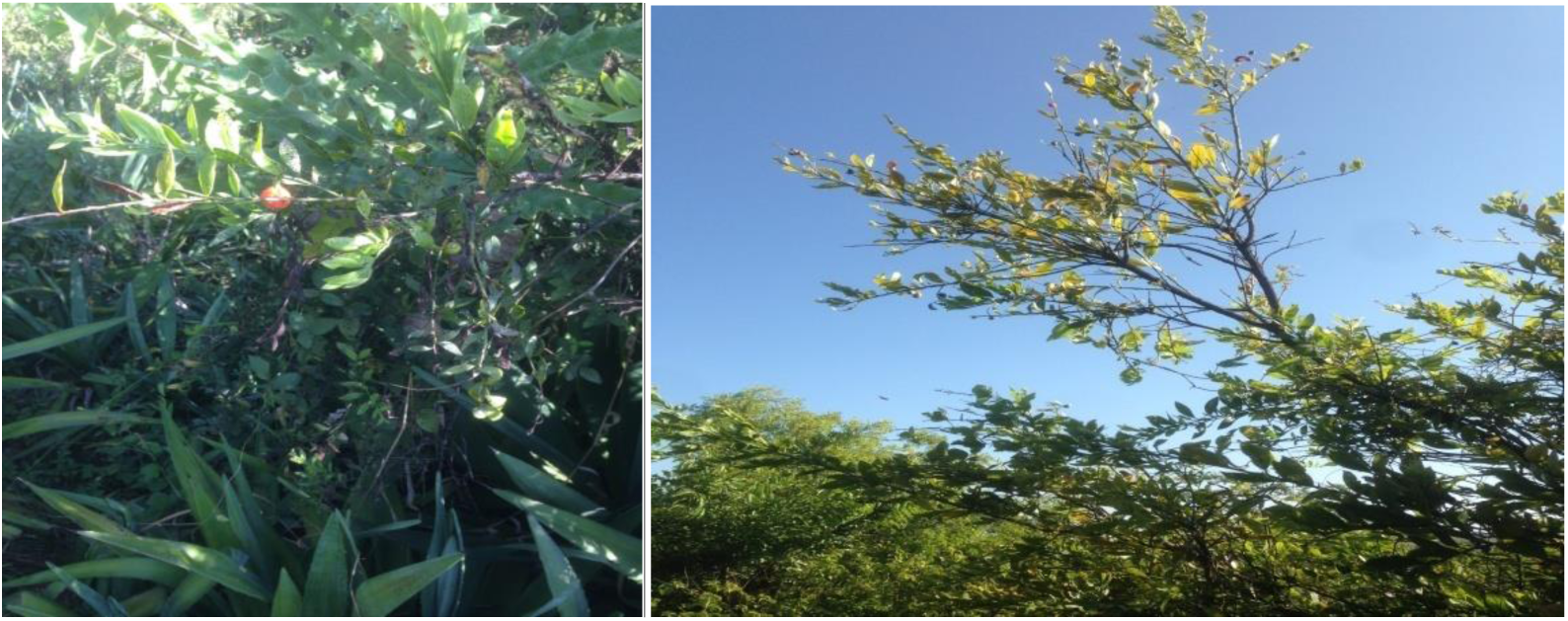
Osyris Lanceoleta in Bazawit Hill.

### 2.2. Experimental design and treatments

Three factors in relation to new leaf intiated success were investigated.The effect of the season at which cuttings were collected in Novmber, the effect of origin of the stem cutting within a shoot portions (i.e. terminal); and the effect of rooting hormone (lndole-3-Butyric Acid) application at concentrations of 0, 50, 100 and 150 ppm. The control set (0 ppm) was treated with distilled water.

For stem cuttings, the terminal portion was taken from the tip of the shoot down to 15 cm, it is ensuring that each stem cutting contained at least two nodes. Auxin concentration was applied by dipping the trimming ends of the cuttings (2-3 cm) into IBA solution for 8 hrs. The IBA concentrations were prepared by dissolving IBA powder diluting with distilled water to make a desirable concentration. Each treatment was replicated four times and for controls eight replication and making a total of 20 for the whole experiment.

**Figure 2.**
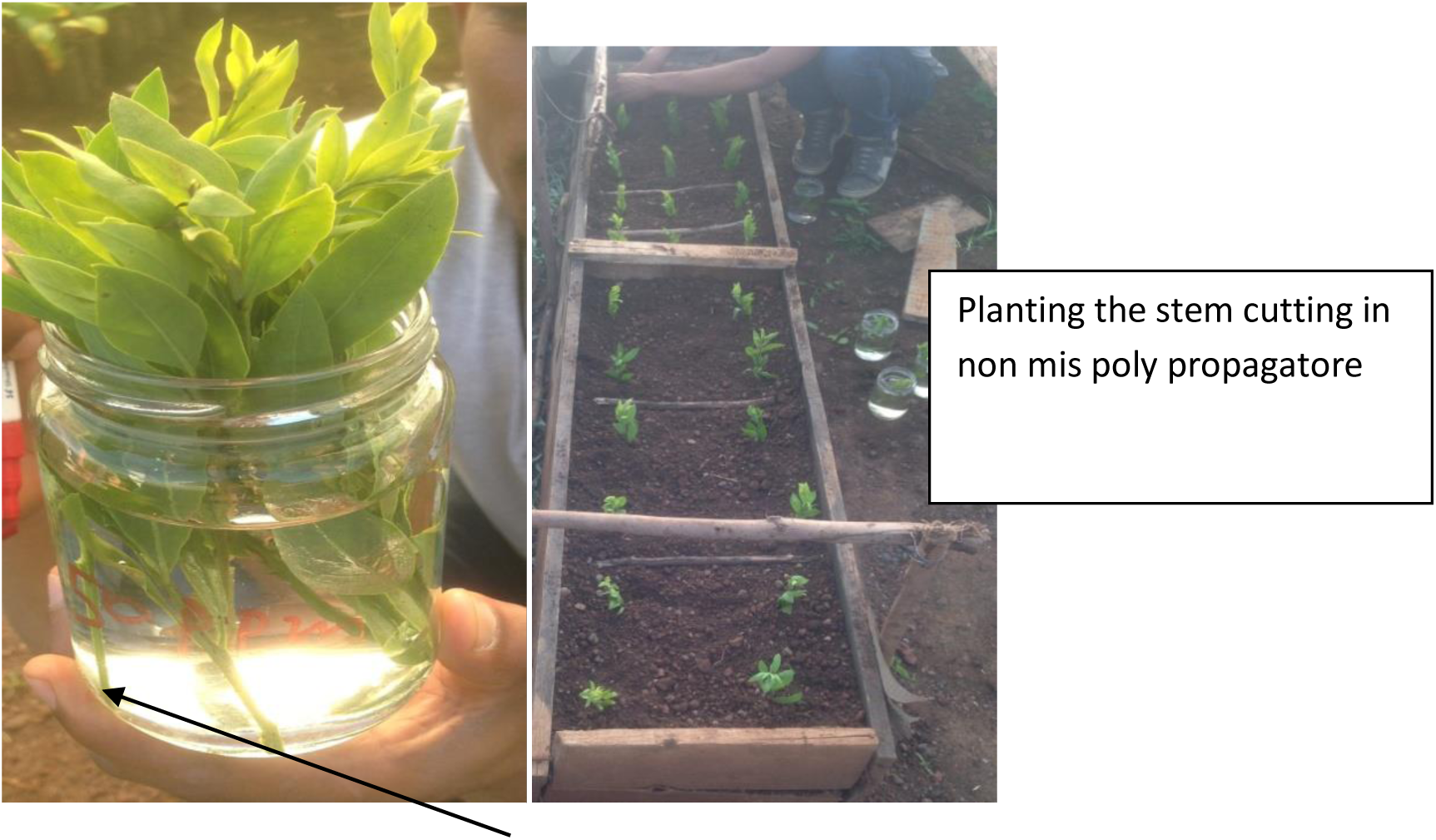
Inserting stem cutting in IBA horomone solution.

### 2.3. Experimental establishment and management

Stem cuttings were collected from sprouts of the mature trees at Bzawit hill. Cuttings were collected in the morning and transported to the experimental site in Bahrdar tissue culture laboratory where further processing and preparation took place. Before planting, stem cuttings were treated with IBA concentrations by dipping the trimming ends for 8 hrs by 0 ppm,50 ppm,100 ppm and 150 ppm of IBA concentartion and planted in a none mist poly-propagtore. Cuttings were initially raised new buds after 5 weeks from planting date. Non mist propagator (Length=196 cm, width =37 cm and hight=15cm) were filled with sand(30%), redish soil(25%) and forest soil(45%).The propagatore were perforated at the bottom to allow easy drainage of water and covered by white plastic. Watering of the cuttings was done every days in the after none and after 6 weeks of the experment 50 % of the treatment formed new leaf and the untreated were 83.3% sucssed.

**Figure 2.**
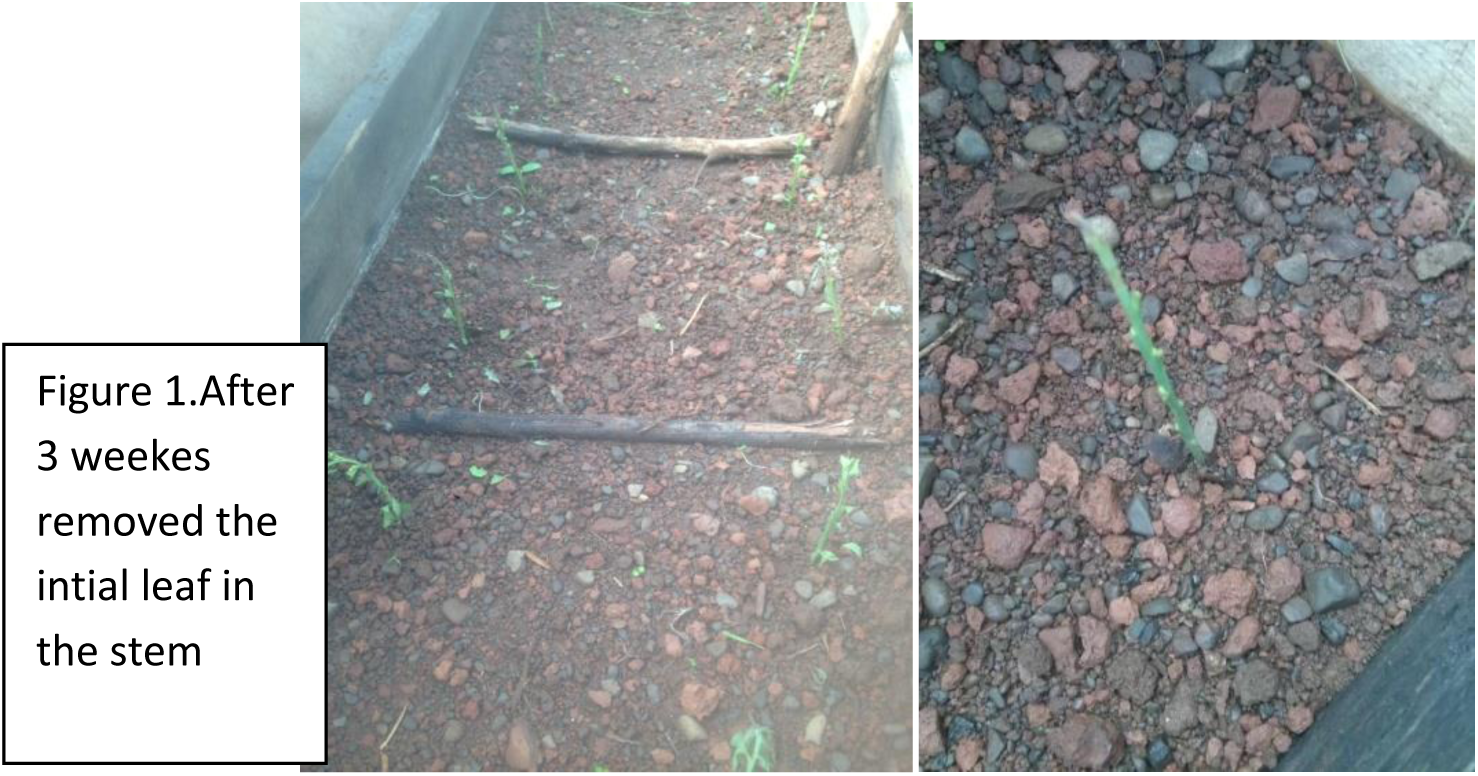
New buds in a non mist poly propagatore in stem cutting after 5 weeks of planting.

**Figure 2.**
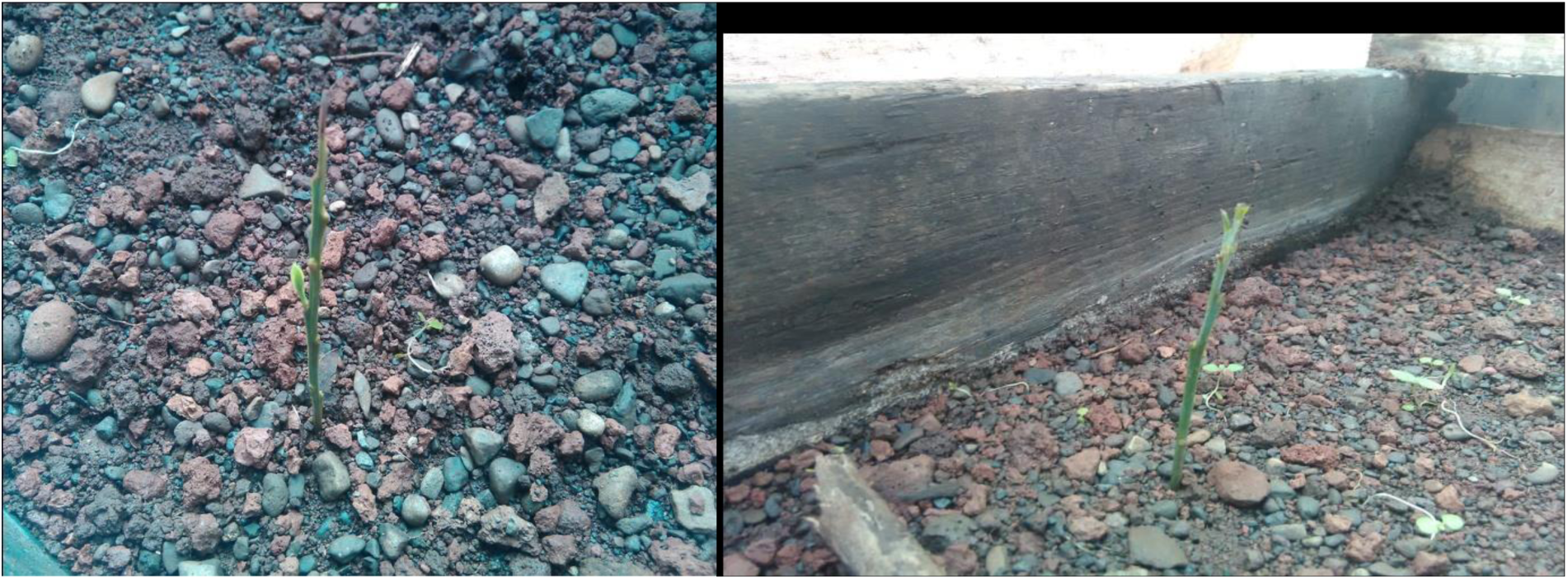
New leaf of Osyris Lanceoleta in the stem in Non mist polypropagatore after 6 weeks.

## 4. Results

New leaf success (the number of cuttings that leafed) was achieved in stem cuttings that were collected in Novmbere, 2019. The number of cuttings not differed between hormone concentration and 83.3 % of the un treatead by horon best new leaf formation.

**Table 1.** The effectes of stem cuting treated by IBA hormone and Distiled water.

| IBA concentration | Numbers of stem cuting | Numbers of responed<br>new shots after 6 weeks | New shots<br>fromation in % |
| --- | --- | --- | --- |
| 50 ppm | 4 | 2 | 50% |
| 100 ppm | 4 | 2 | 50% |
| 150 ppm | 4 | 2 | 50% |
| 0ppm(Distiled<br>water) | 6 | 5 | 83.3% |

## 5. Conclusion

Due to Osyris Lanceoleta seed is very limited and sensitive to fungi, it is very difficult to propagate by seed. However, this research finding via using the advances of plant propagation method it provide new options for conserving and multiplication of Osyris lanceolata species using IBA hormone at different concentration and distiled water by taking stem plant material of Osyris Lanceoleta in non -mist poly propagatore.

## 6. Acknowledgments

Ethiopian Enviroment and Forest Research Institute appreciates highly Ethiopian Enviroment, Forest and Climate Change Commision for fully funding this research via UNDP Institutional Strengthening for the Forest Sector Development Program.

